# Rab7 mediates Wnt-dependent mesoderm patterning and gastrulation in *Xenopus*

**DOI:** 10.1101/2020.08.14.251298

**Authors:** Jennifer Kreis, Fee M. Wielath, Philipp Vick

## Abstract

Early embryogenesis requires tightly controlled temporal and spatial activity of growth factors. Modulation is achieved at multiple levels, from cellular transcription to tissue-scale propagation, concerning activation, maintenance, or termination of signaling. Intracellularly, the endolysosomal system emerged as important factor for such regulations. Endocytosis and early endosomes mainly orchestrate activation and transduction of signals, while late endosomes are thought to mediate lysosome-based termination. Only few reports attributed activating roles to late endosomes, mainly for Wnt pathway activation. In *Xenopus*, few is known about developmental roles of endosomal regulators, or their function for signaling, especially true for late endosomes. Therefore, we chose to analyze a hypothesized positive role of Rab7 in this context, a small GTPase mainly known for its role as a late endosomal regulator. We analyzed if Rab7 participates in early activation of growth factor signaling, focusing on Wnt signaling. We found *rab7* expressed very localized throughout development, and inhibition resulted in strong gastrulation defects. Concomitantly, Rab7 was strictly required for both, exogenous activation and endogenous Wnt-mediated patterning of the mesoderm.

**Summary Statement:** The late endosomal regulator Rab7 is required for gastrulation movements and axis elongation in *Xenopus* by positively regulating Wnt signaling-mediated mesoderm patterning.

## Introduction

Early embryonic processes like germ layer formation, induction of body axes, neural induction or tissue differentiation require tightly controlled temporal and spatial activation of specific sets of signaling pathways. Regulation of endocytosis or membrane trafficking can control activation, intensity, or duration of intracellular signal transduction upon ligand-mediated activation of transmembrane receptors (Sigismund et al., 2012). However, this has only been analyzed in few developmental processes *in vivo*, as altering such basic cellular processes can have dramatic effects.

Classically, endocytosis of plasma membrane localized receptors is considered a way of downregulation of signaling pathways. Receptors are translocated to early endosomes (EE) by endocytic vesicles, which quickly fuse with EE membranes, representing first intracellular sorting platforms (Platta and Stenmark, 2011). From here, activated receptor complexes can be inactivated (e.g. ligand-separated), and recycled back to the plasma membrane, which enables repeated pathway activation. Alternatively, such activated receptors are retained in EE membranes while these organelles mature into late endosomes (LE). Here, transmembrane cargo can be translocated into its intraluminal vesicles (ILV) by successive inward budding of the limiting membrane, a process characteristic for maturing LE, and which is performed by the ESCRT (endosomal sorting complexes required for transport) complex (Hanson and Cashikar, 2012). When translocated into ILV, receptors are separated from the cytoplasm, and this is generally thought to be a one-way route to degradation for any cargo, as matured LE fuse with lysosomes for acidic degradation of contents. Thus, in most cases analyzed so far, LE represent an intermediate step between EE and degradation, with no return once LE are matured and transmembrane receptors are translocated into ILV (Dobrowolski and De Robertis, 2011; Horner et al., 2018; Katzmann et al., 2002).

While degradation and recycling as ways of modulating signal strength are well established, the role of endocytosis and endosomes for activation or maintenance of signaling pathways are much less understood. It is known that TGF-β (transforming growth factor beta), BMP (bone morphogenetic protein), Notch, and both, canonical and non-canonical Wnt signaling pathways require endocytosis to activate intracellular signaling upon ligand binding. In addition, BMP and TGF-β need to be transported to EE, where specific adapters are located, which are in turn required to bridge the receptor kinase to its substrate, a Smad family transcription factor (Brunt and Scholpp, 2018; Butler and Wallingford, 2017; Dobrowolski and De Robertis, 2011; Fürthauer and González-Gaitán, 2009; Platta and Stenmark, 2011). A rarer case is the description of a positive role of LE for pathway activation – as opposed to a simple role for degradation. This has been suggested to be the case for EGF (epithelial growth factor) receptor-mediated MAPK (mitogen-activated protein kinase) activation under certain circumstances (Platta and Stenmark, 2011; Teis et al., 2002). Further, for canonical Wnt signaling (from here on simply ‘Wnt signaling’), it has been demonstrated that LE are indispensable for sustained pathway activation, i.e. for maintaining signaling after endocytosis-mediated activation of the Wnt receptor complex (Platta and Stenmark, 2011; Taelman et al., 2010; Vinyoles et al., 2014).

Rab family proteins are a group of small GTPases that regulate most membrane trafficking processes by transiently binding membranes with their C-terminal prenyl anchor. Each Rab is specific to certain types of membranes/organelles and orchestrates the recruitment of a specific set of effectors, thereby giving the membrane a specific ‘identity’ and function. For most routes of cellular membrane trafficking, one or few Rab proteins serve as process-specific molecular switches (Stenmark, 2009). Although, as judged by their important general roles in cellular transport, many Rab proteins are categorized as ‘housekeeping genes’, they might also be involved in specific processes, like regulation of signaling pathways. An example is the endocytic/EE regulator Rab5, which is often required for internalization of receptor complexes and can therefore regulate intracellular output of signaling.

The LE regulator Rab7a (from here on simply Rab7), and its low expressed, tissue-specific paralog Rab7b are mainly found on LE. Initially, they are recruited to endosomal membranes by EE-located Rab5, which is then replaced by Rab7 when these structures mature into LE. Thus, Rab7 is often used as organelle marker for LE, and more importantly, as a recruiter of many effectors, it is also the main regulator of LE maturation and function (Huotari and Helenius, 2011; Stenmark, 2009). Concerning signaling pathways, it is thus considered as a permissive regulator of endolysosomal degradation, i.e. required for membrane receptor degradation and termination of signaling (Platta and Stenmark, 2011). While this is straightforward logic, controlling the Rab5 to Rab7 conversion, or Rab7 activity itself, could also be a way of positively regulating activity of downstream signal transducers. For instance, this might be the case for canonical Wnt signaling, where fully functional LE have been shown to be required for sustained target gene activation. In addition, the canonical Wnt pathway has been reported to be able to positively influence expression of endosomal regulators, representing an example of a direct influence on the endolysosomal system in a positive feedback loop (Ploper and De Robertis, 2015; Ploper et al., 2015; Taelman et al., 2010).

In fact, studies dealing with the *in vivo* function of Rab7 are rare, especially in a developmental context, and most information about its influence on pathway activity derives from work in cell culture, i.e. from out-of-tissue contexts (Guerra and Bucci, 2016). This might be due to the important role of Rab7 in every cell, causing classical knockout (KO) approaches to result in embryonic lethality. This is exemplified by work in the mouse, where *rab7* KO resulted in early death before embryonic day 8. KO embryos had strong defects in endosomal transport, especially in the anterior visceral endoderm (AVE), which resulted in antero-posterior (AP) patterning and AVE migration defects, and thus in failure to complete gastrulation (Kawamura et al., 2012; Stower and Srinivas, 2014). If these phenotypes were related to mis-regulation of Wnt signaling or other developmentally important pathways was not addressed in this study. So, while cytoskeletal or endosomal regulators like Rab proteins turned out to be important modulators of pathway activity at multiple levels, not many studies analyzed this in a developmental context.

In this work, we chose to analyze the *in vivo* function of *rab7* and its potential participation in Wnt pathway activation during early development of *Xenopus laevis*. In contrast to its housekeeping role, we found *rab7*-mRNA specifically enriched in distinct types of tissues during early development, reflecting dynamic changes of enhanced requirement. Using morpholino-mediated knockdown and CRISPR/Cas9-mediated genome editing, we found that loss of *rab7* resulted in gastrulation defects without impacting embryonic organizer induction on the dorsal side. Further, *rab7* was required for ligand-mediated activation of exogenously induced, as well as for endogenous canonical Wnt signaling in the mesoderm.

## Results

### Loss of *rab7* results in gastrulation defects

We speculated *rab7* could show distinct spatial enrichment of mRNA expression during early *Xenopus* development. If the case, such enhanced abundance would give indications about tissue-and process-specific requirements of *rab7*. Indeed, expression analysis by *in situ* hybridization (ISH) revealed very dynamic spatial and temporal signals. Strong maternal expression was found in the animal half of cleavage stages, a signal still detected at the onset of zygotic transcription after midblastula transition (MBT; Fig. 1A,B; Fig. S1A). At early gastrula stages, transcripts were mainly found in the deep mesodermal ring, omitting the dorsal lip, i.e. the anterior/head part of Spemann-Mangold organizer (Fig. 1C,D; Fig. S1B). During late gastrulation, stronger signals were detected in the neural plate ectoderm and in the axial, notochordal mesoderm, latter of which continued to be positive for *rab7* during neurulation (Fig. 1E; Fig. S1C). By then, ectodermal expression became more restricted to the lateral neural plate, and later in the deep cells of the neural tube and brain tissues (Fig. 1F-H; Fig. S1D-E). In following tailbud stages, *rab7* signals were detected in the cement gland and dermal areas, slightly in the notochord, and strong in the pronephric kidney, eyes, ventro-lateral neural tube, pharyngeal arches, trigeminal nerve complex, dorsal fin mesenchyme, and in the trunk neural crest cells (Fig. 1I,J; Fig. S1F,G; and data not shown). This analysis supported our hypothesis that *rab7* could be required for early embryonic development by participating in regulation of signaling activity in multiple tissues.

**Fig. 1.**
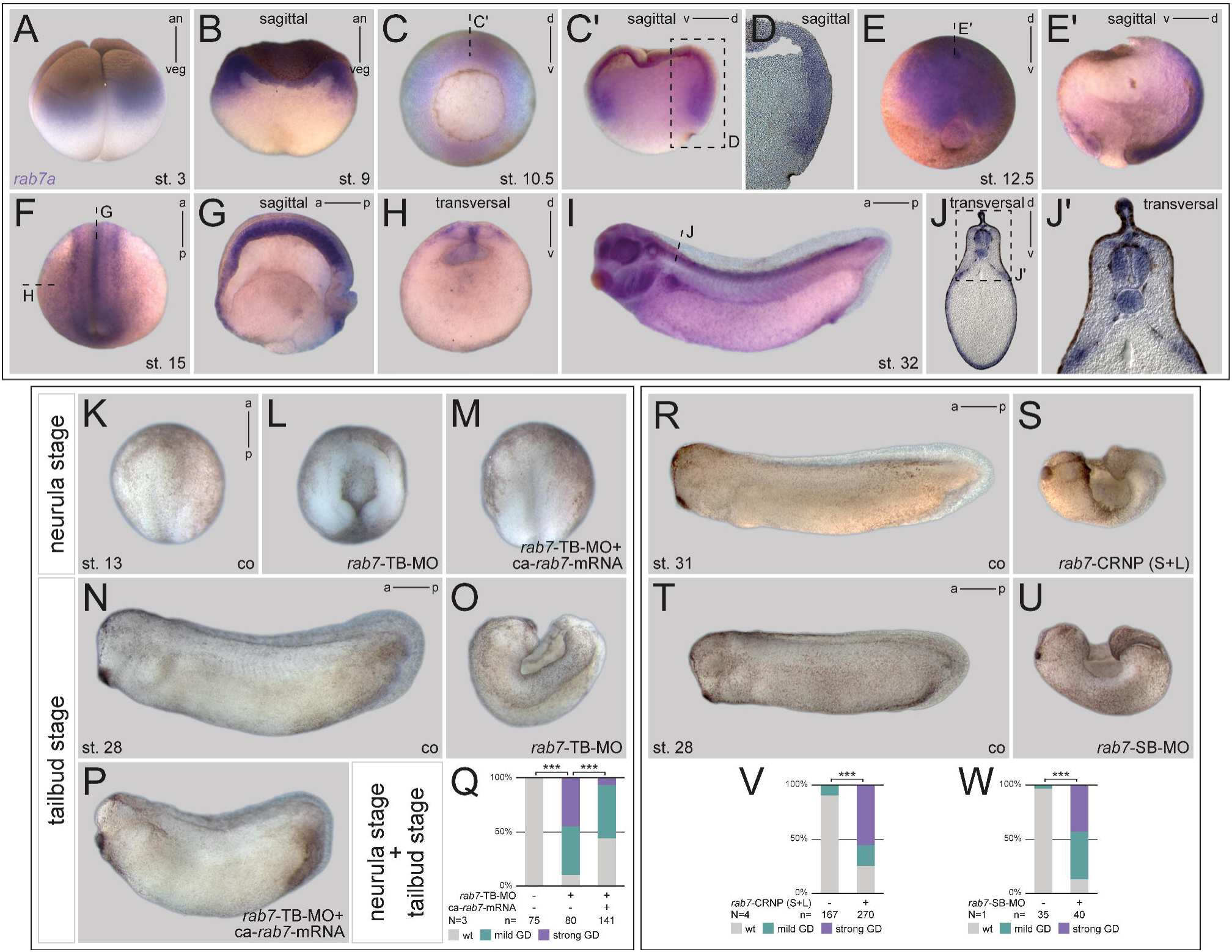
*rab7* shows a dynamic expression and inhibition resulted in gastrulation defects. (A, B) Expression of *rab7* mRNA in animal hemisphere at st. 3 and sagittal section of late blastula stage. (C) Upon gastrulation transcripts were enriched in deep mesodermal ring, (C’) sagittal section indicated in (C), (D) blow-up of (C’). (E) At st. 12.5 *rab7* accumulates in neural plate ectoderm; (E’) sagittal section of (E). (F) During neurulation transcripts get restricted to notochord, neural tube and brain tissue, (G) sagittal and (H) transversal section indicated in (F). (I) In tailbud stages transcripts were detected in the notochord and pronephric and head tissues, (J) transversal section indicated in (I), (J’) blow-up of (J). (K,N) Control specimen at st. 13 and 28. (L,O) Injection of *rab7*-TB-MO in dorsal lineage caused gastrulation defects, resulting in severe dorsal phenotypes. (M,P) Co-injection of *ca-rab7* mRNA rescued loss of function phenotype of morphant embryos. (Q) Quantification of results in (K-P). (R,T) Tailbud control embryos; (S,U) siblings treated with *rab7*-CRNP (S+L) or *rab7*-SB-MO showing dorsal phenotypes. (V,W) Quantification of results in (R,S and T,U). a, anterior; an, animal; ca, constitutive active; co, control; CRNP, Cas9 Ribonucleoprotein; d, dorsal; GD, gastrulation defect; p, posterior; SB-MO; st., stage; v, ventral; veg, vegetal; wt, wildtype.

Next, we wanted to test such an early *in vivo* requirement of *rab7* using a loss-of-function approach. We designed a morpholino oligomer (MO) targeting the 5’UTR of the L- and S-form of *Xenopus laevis rab7* to block translation initiation of both homeologs (*rab7*-TB-MO). Morphant embryos passed through cleavage and blastula stages without detectable phenotypes (not shown). However, subsequent gastrulation movements were strongly inhibited, causing failure to close the dorsal blastopore and resembling strong convergent extension (CE) defects in about 50% of the embryos at early neurula stage (Fig. 1K,L). This phenotype became more prevalent at tailbud stages, with further extension of the AP axis in control specimens, while morphant embryos remained wide open dorsally with a strong dorsal curvature (Fig. 1N,O). Importantly, co-injection of a constitutively-active (ca) *rab7* mRNA was able to rescue the strong gastrulation and CE phenotype in a highly significant manner, demonstrating specificity of the observed MO effect (Fig. 1M,Q). At tailbud stages, nearly all rescued embryos were able to close the dorsal blastopore and to elongate their AP axis, albeit with a slight delay in AP elongation (Fig. 1P,Q). To further underline specificity of this effect we next designed single guide RNAs (sgRNA) to target the genomic loci of both *rab7* homeologs for CRISPR/Cas9-mediated mutagenesis, either in parallel, or individually. KO efficiency of injected embryos was determined by sequencing and subsequent analysis of indel distribution (Synthego ICE; for details see Method section). These analyses resulted in a predicted gene editing rate between 60% and 88% for L- or S-forms of the different sgRNAs (Fig. S1H-J). Genome editing of *rab7* L/S at the one-cell stage caused strong gastrulation defects, again resulting in tailbud stages with open dorsal part in over 50% of specimen, resembling the phenotype shown for morphants (Fig. 1R,S,V). Interestingly, while selective targeting of homeolog L with sgRNA (L) caused also a similar phenotype in about 25% of injected specimen (Fig. S1K-M), no gastrulation defects were observed by only targeting homeolog S with sgRNA (S) (not shown). Finally, we also designed a splice-blocking (SB) MO targeting the splice acceptor site at intron 2 of the *rab7* pre-mRNA to prevent splicing, and thus should cause translational read-through and early termination (Fig. S1N). Successful inhibition of splicing could be demonstrated by RT-PCR for both homeologs, resulting in intron retention for each form (Fig. S1O). Additionally, injection of the *rab7*-SB-MO resulted in significant reduction of *rab7*-transcript amounts in morphant neurula or tailbud stage embryos, indicating nonsense-mediated decay of unspliced *rab7*, and thus successful knockdown of zygotic transcripts (Fig. S1P-S). Phenotypically, *rab7*-SB-MO injected embryos showed also gastrulation defects, but to a lower extend and degree (Fig. 1T,U,W). Those specimens who managed to close the blastopore were raised until tailbud stages. Such milder affected morphants displayed deficits in AP elongation, indicating a milder CE phenotype (Fig. S1V,W). Finally, by raising the few surviving *rab7*-TB-MO, or sgRNA (S+L) injected embryos, this late phenotype could be phenocopied in early tadpole stages, again showing specificity of the effect (Fig. S1T,U,X,Y). In summary, our loss-of-function approach demonstrated a requirement of *rab7* for early embryonic development.

### Rab7 is required for convergent extension and notochord morphogenesis

To better understand this gastrulation phenotype, we specifically knocked down *rab7* in the dorsal mesoderm. When mid-sagittal sections were analyzed at early gastrula stages, *rab7* morphants formed a lip, but involution movements and archenteron formation were significantly impaired (Fig. 2A-D). Interestingly, bisections revealed a concomitant lack of Brachet’s cleft, implying incorrect tissue remodeling during early gastrulation (Fig. 2C-F; (Fagotto, 2020). These phenotypes became also apparent when such embryos were analyzed for notochordal *noggin* (*nog*) expression at mid-gastrula, supporting strongly impaired axial mesoderm elongation and reduced *nog* expression itself, the latter one indicating a failure in maintaining notochordal fates in morphant tissue (Fig. 2G-J). Analysis of cell-matrix/cell-cell adhesion revealed a reduction of β1-Integrin and Ctnnb1 (β-catenin) in the axial mesoderm of morphants (Fig. 2K-M and Fig. S2A-C). Similar reductions were observed in animal caps, where loss of *rab7* reduced Ctnnb1 in targeted areas, suggesting a general effect (Fig. S2D).

**Fig. 2.**
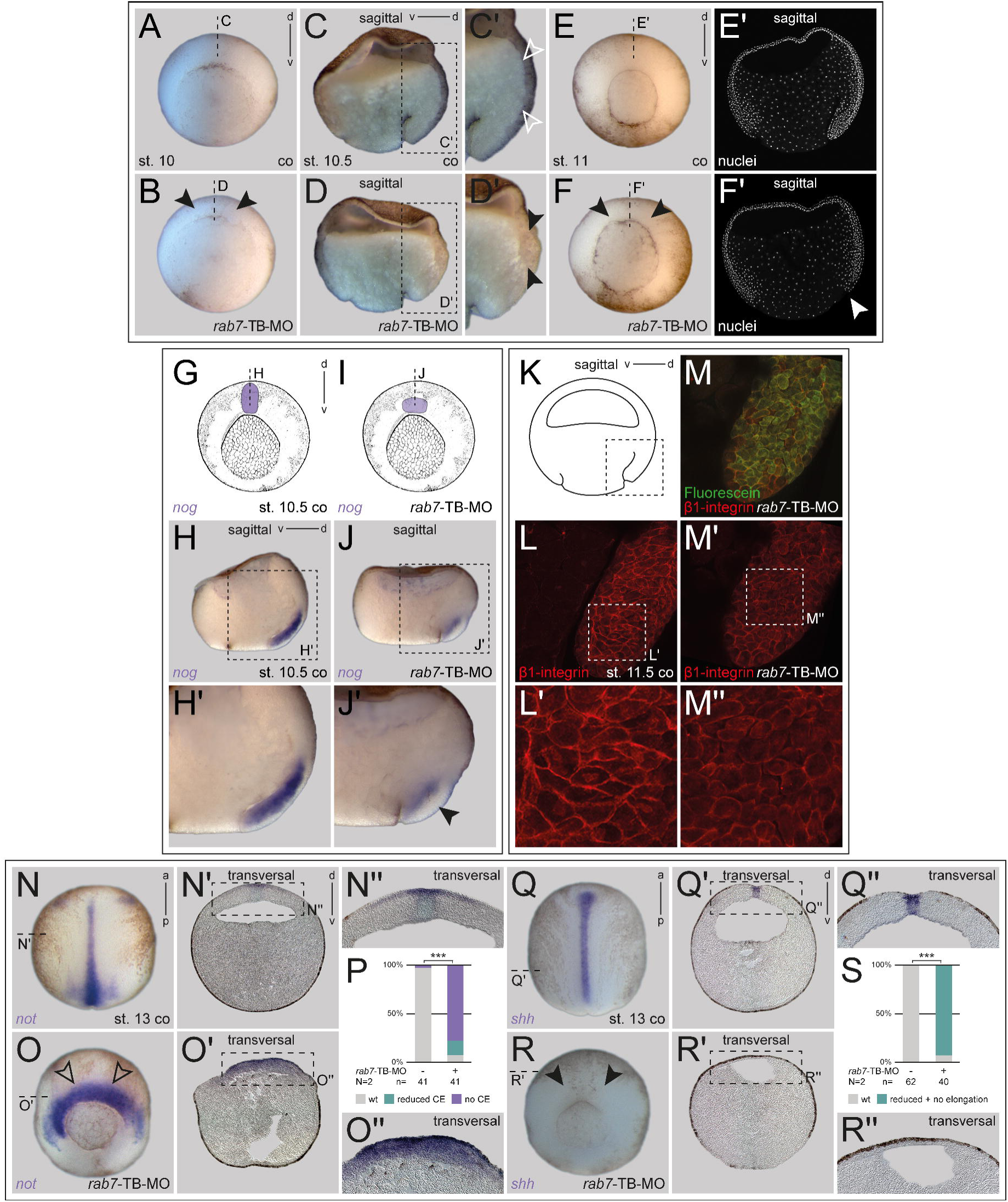
*rab7*-deficient embryos show convergent extension defects and impaired notochord morphogenesis. (A) Untreated embryos at the begin of gastrulation forming a dorsal lip, (B) which can also be observed in *rab7* deficient siblings (black arrowheads). (C) In st. 10.5, sagittal sectioned control embryos reveal early involution movements, in blow-ups (C’) formation of Brachet’s cleft was visible (white outlined arrowheads); (D) dorsal *rab7* knockdown specimen rarely showed involution in sagittal sections or (D’) Brachet’s cleft formation in blow-ups (black arrowheads). (E) Older control embryo with evenly shaped lip around the blastopore; (E’) sagittal section with Hoechst stained nuclei revealed proceeded dorsal involution. (F) Morphant siblings developed an irregular shaped lip around the blastopore (black arrowheads), and (F’) did not show any involuting tissue in Hoechst stained sagittal section (white arrowhead). (G) Schematic figures indicate wildtype *nog* expression of control embryos and (I) reduced expression of *nog* and concomitant failed notochord elongation of dorsal *rab7*-TB-MO injection. (H) Sagittal section and (H’) blow-up exhibited proceeded notochord elongation marked by *nog* in untreated embryos. (J) Sagittal section and (J’) blow-up clearly depicted impaired *nog* expression (black arrowhead). (K) Schematic figure of sagittaly sectioned mid-gastrula indicates detailed view of dorsal lip shown in (L-M). (L,M) sections stained for β1-integrin (red) and fluorescein dextran (FD) as lineage tracer (green). (L) Normal distributed β1-integrin at cell surface of wildtype dorsal lips, (L’) blow-up of (L) showing normal cell shape. (M) FD staining indicates cells positive for *rab7*-TB-MO, (M’) morphant cells reveal reduced β1-integrin levels and (M’’) blow-up of (M’) exhibit an altered cell shape. (N,Q) Early wildtype neurula embryos showing elongated notochords marked by the expression of *not* or *shh*. (N’,Q’) Transversal sections indicated in (N,Q) revealed *not* and *shh* expression in the axial neural plate and *shh* additionally throughout the whole notochord. (N’’,Q’’) blow-ups of (N’,Q’). (O,R) Dorsal *rab7*-TB-MO injection resulted in embryos failing to elongate their notochord, although *not* expression intensity (black outlined arrowheads) was not reduced in comparison to *shh* (black arrowheads). (O’) Transversal section of morphant embryos showed lateral expanded expression of *not* above the dorsal lip and (R’) severely reduced *shh* expression. (O’’,R’’) blow-ups of (O’,R’). (P,S) Quantification of results in (N,O and Q,R). *Abbreviations as indicated in Fig. 1*

As these phenotypes also indicated a failure in CE and thus, to form a proper elongated notochord subsequently, we next examined notochord fate and appearance directly. When we checked expression of the marker *notochord homeobox* (*not*) in morphant embryos at early neurulation, a lack of CE was obvious, explaining the embryos’ inability to close the dorsal part of the blastopore (Fig. 2N-P). When using the *rab7*-SB-MO for dorsal knockdown, a milder but significant effect was observed as well, (Fig. S2E-G). Interestingly, while *not* expression was not reduced in the axial mesoderm and extended into the lateral somitic area, analysis of *sonic hedgehog* (*shh*) in the same set of experiments revealed a different effect. Morphant embryos showed a similar lack of CE but expression intensity of *shh* was reduced in most cases, again suggesting partial lack of notochordal fate (Fig. 2Q-S).

Morphant embryos were grown to tailbud stages to analyze notochordal tissue differentiation. In these stages, *not* expression was also not reduced, but appeared enhanced in milder affected specimens, especially in the mid-trunk area, where in wildtype embryos expression had already faded by that stage (Fig. S2H-J). Overall, notochordal appearance in sagittal sections confirmed attenuated CE. Stronger affected specimens developed open dorsal tissues and mostly retained *not* expression, but often split as two separated areas relocated medially (Fig. S2K). Staining such embryos with an MZ-15 antibody, which detects outer keratan sulphates of the notochordal sheet, revealed absent or strongly reduced signals, supporting a lack of notochord differentiation (Fig. S2L-O). These phenotypic analyses showed that *rab7* is participating in axial tissue morphogenesis and CE-dependent notochord formation during gastrulation.

### Rab7 is required for dorsal mesoderm specification but not for the organizer

To understand the role of *rab7* in this context, we next analyzed if organizer induction was impaired by targeting this dorsal lineage. Neither knockdown with *rab7*-TB-MO or *rab7*-SB-MO, nor CRISPR-induced KO impacted on correct organizer induction, as judged by unaltered expression patterns of the organizer genes *goosecoid* (*gsc*) and *chordin* (*chd*; Fig. 3A-C and Fig. S3A-H). Next, we wanted to test if early pre-gastrula dorso-ventral (DV) axis formation was altered in *rab7* morphants. We knocked down *rab7* and checked dorsal *chd* or ventral *ventx1* expression to analyze for altered Bone Morphogenetic Protein (BMP) gradient. Both genes were found in similarly extending domains as in control embryos (Fig. 3D-F and Fig. S3I-K). This suggested that *rab7* is dispensable for both, organizer induction and DV patterning.

**Fig. 3.**
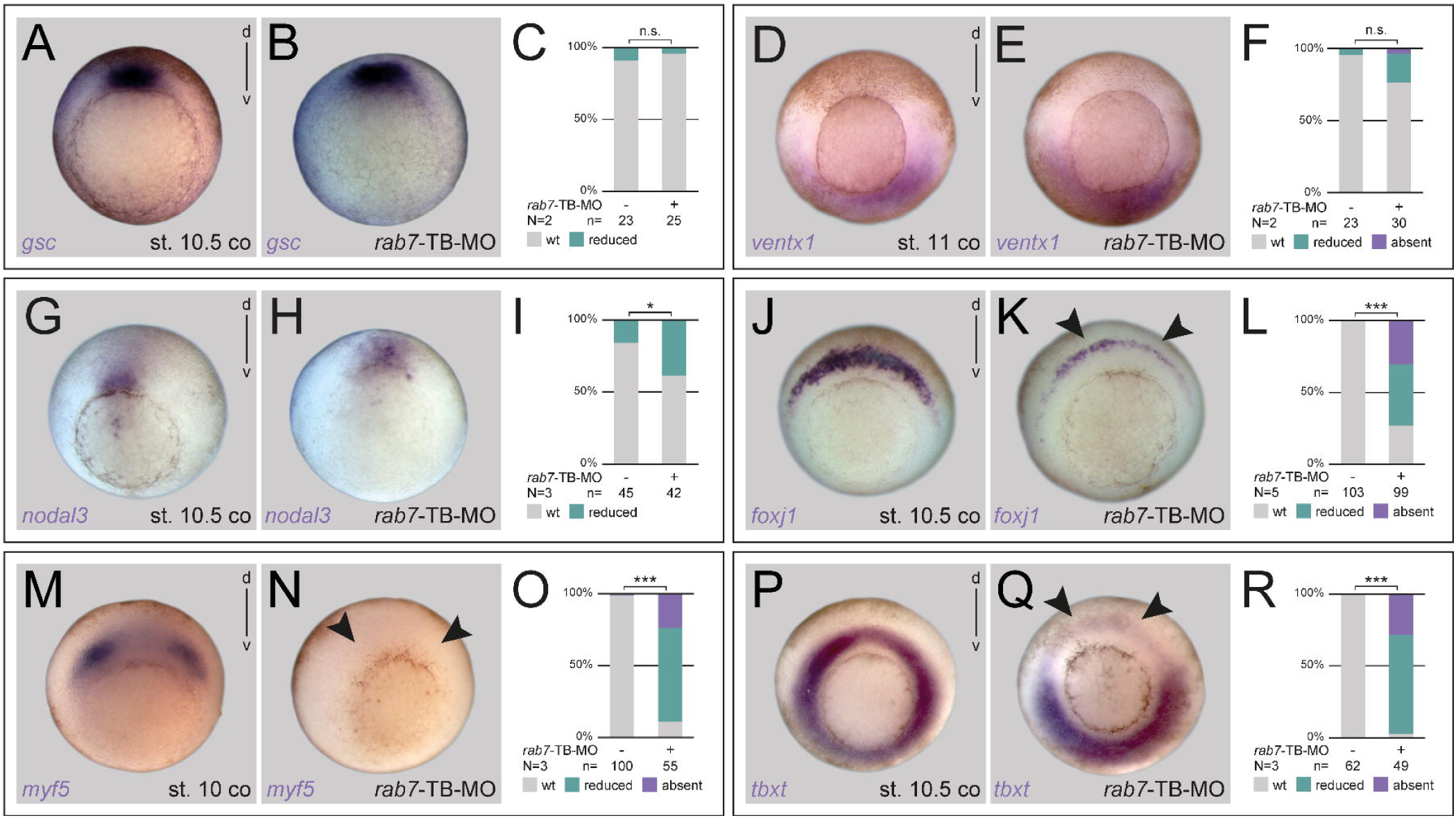
Dorsal mesoderm specification requires Rab7 independent of the organizer. (A,D,G,J,M,P) Untreated control specimen with wildtype *gsc, ventx1, nodal3, foxj1, myf5* and *tbxt* expression, respectively. (B,E) *rab7* loss of function in dorsal or ventral lineage did not alter *gsc* or *ventx1* expression, respectively. (H) *nodal3* expression was slightly reduced in some *rab7* morphant embryos. (K,N,Q) expression of *foxj1, myf5* and *tbxt* were severely affected in specimen with *rab7* deficiency (black arrowheads). (C,F,I,L,O,R) Quantification of results in (A-B,D-E,G-H,J-K,M-N or P-Q). *Abbreviations as indicated in Fig. 1*

As a main regulator of LE, *rab7* would be expected to be indispensable for maternally activated canonical Wnt signaling, and thus for endogenous organizer induction. However, the reduced expression pattern of *shh* suggested that *rab7* might be required for other early dorsal mesoderm genes as well. Therefore, we checked expression patterns of *nodal3* and *forkhead box J1* (*foxj1*), two Wnt-dependent marker genes expressed in the superficial mesoderm (SM), i.e. the outer layer of the trunk organizer tissue (Glinka et al., 1996; Smith et al., 1995; Stubbs et al., 2008; Walentek et al., 2013). Interestingly, while the very early expressed *nodal3* was reduced in a fraction of embryos, *foxj1* was strongly reduced after loss of *rab7* (Fig. 3G-L). The paraxial presomitic expressed marker *myogenic factor 5* (*myf5*), also well-known to be dependent on Wnt (Kjolby and Harland, 2017; Shi et al., 2002), was strongly downregulated (Fig. 3M-O), indicating a potential fate shift as indicated by the lateral extension of *not* into these areas (cf. Fig. 2O). Finally, we wanted to test, if mesodermal identity was altered more generally, as this has been reported to depend on Wnt pathway activation as well (Vonica and Gumbiner, 2002). Analysis of *T-box transcription factor T* (*tbxt*, also known as *brachyury*) revealed a significant reduction of expression after *rab7* knockdown in this area (Fig. 3P-R), explaining the morphogenetic phenotype in the axial mesoderm (cf. Fig. 2). The down-regulation of these marker genes suggested a role of *rab7* for Wnt-dependent development of the dorsal mesoderm, probably downstream or in parallel of endogenous organizer induction.

### Rab7 is required for canonical Wnt pathway activation *in vivo*

In order to support our conclusion that the loss of *rab7* blocked an endogenous dorsal Wnt signal, and to bypass the possibility that maternally deposited *rab7* mRNA or protein would ‘cover’ its requirement for Wnt-induced organizer induction in our experiments, we next used radial injections of *wnt8a* mRNA, which is well-known to dorsalize the embryo (Hikasa and Sokol, 2013; Smith and Harland, 1991). Injections caused radial expression of dorsal-specific organizer genes and erased that of ventral-specific *ventx1* (Fig. 4A,B,D and Fig. S4E-H). Importantly, co-injection of *rab7*-TB-MO restored the DV-axis highly significantly without impacting on endogenous organizer-specific expression of *chd* or *nog* (Fig. 4C,D; Fig. S4A-D). Early gastrula embryos tested for *ventx1* showed restoration of their DV axis but *ventx1* expression was partially reduced (Fig. S4G,H). These results suggest that *rab7* is required for ligand-dependent activation of canonical Wnt signaling *in vivo*.

**Fig. 4.**
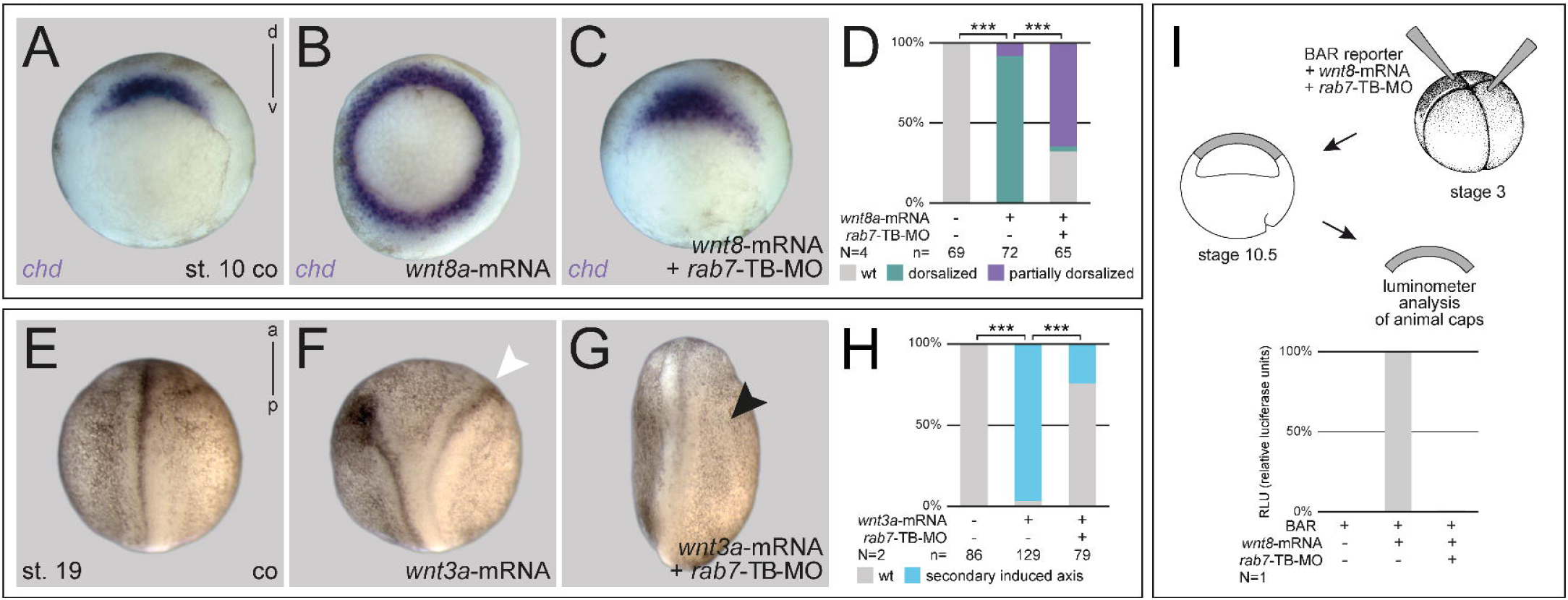
Rab7 is required for canonical Wnt pathway activation *in vivo*. (A) Wildtype dorsal expression of *chd* in comparison to specimen with (B) radial injected *wnt8a*-mRNA showing an extended *chd* expression domain around the blastopore. (C) Co-injection of *rab7*-TB-MO restricted additional *chd* expression to normal wildtype state. (D) Quantification of results in (A-C). (E) Untreated control specimen, (F) in comparison to induced double axis (white arrowhead) after ventral *wnt3a*-mRNA injection; (G) parallel injection of rab7-TB-MO inhibited secondary axis formation (black arrowhead). (H) Quantification of results in (E-G). (I) Luciferase-based Wnt reporter assay analysis at st. 10 illustrated that *wnt8a* induced reporter activity was blocked by *rab7* loss of function in animal caps. *Abbreviations as indicated in Fig. 1*

If *rab7* participated in Wnt pathway activation *in vivo*, then knockdown should also prevent induction of secondary axes, the classical readout for canonical Wnt pathway in *Xenopus* (Sokol et al., 1991). Indeed, co-injection of *rab7*-TB-MO was sufficient to prevent *wnt3a*-induced double axis induction in a highly significant manner, whereas the efficiency of *ctnnb1* (*β-catenin*) to induce double axes was not altered after *rab7* knockdown (Fig. 4E-H and Fig. S4I-L). To further rule out that this effect was due to an impairment of processes downstream of organizer induction, we also analyzed secondary organizers for expression of *gsc*, which clearly demonstrated a blockage of the exogenous organizer (Fig. S4M-O). This was also supported by the β-catenin activated reporter (BAR; Biechele et al., 2009), which was used to perform a luciferase-based Wnt assay in animal caps. Here, *wnt8a*-induced reporter activation was also blocked by *rab7* inhibition (Fig. 4I). These results supported our conclusion that *rab7* is required to mediate early Wnt signaling in *Xenopus*. Further, this excluded the possibility that its knockdown could impact on processes downstream of *ctnnb1*-dependent target gene activation. Altogether, from this set of experiments we conclude that the small GTPase Rab7 is required for Wnt pathway activation in early frog embryos, upstream of Ctnnb1 stabilization.

### Rab7 is required for specification of the ventro-lateral mesoderm

On the one hand, our experiments have demonstrated that *rab7* participates in the canonical Wnt pathway *in vivo* when ligand-activated exogenously in the mesoderm or ectoderm. On the other hand, endogenous organizer induction was not blocked after loss of *rab7*, yet, dorsal Wnt-dependent mesoderm specification was significantly impaired. Therefore, we next asked if *rab7* also participated in ventro-lateral mesoderm specification, which is known to be dependent on zygotic *wnt8a* (Hoppler and Moon, 1998). During gastrulation, the organizer secretes Wnt antagonists dorsally in the axial mesoderm, while *wnt8a* is expressed in the ventro-lateral mesoderm. Using targeted injections, we blocked *rab7* only in the ventral mesoderm. Morphant embryos developed milder phenotypes with low lethality rates and could thus be raised until tadpole stages. Such embryos showed ventro-posterior malformations, which became more pronounced as early tadpoles, when tail formation was strongly inhibited in most cases (Fig. 5A-C and S5A,B).

**Fig. 5.**
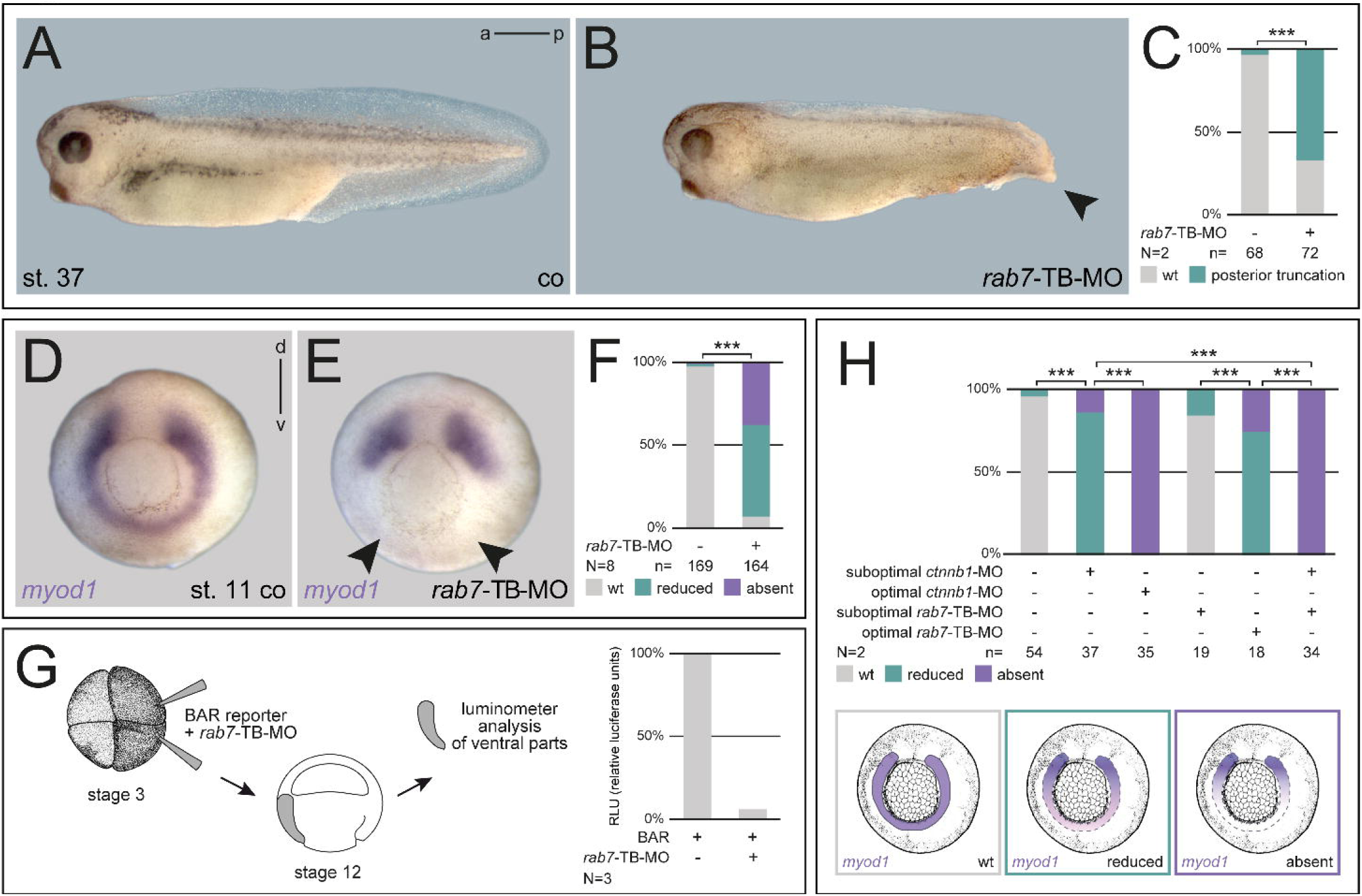
Specification of the ventral mesoderm requires Rab7. (A) Untreated early tadpoles with normal tail development, (B) which was impaired by *rab7* loss of function in ventral lineage resulting in posterior truncations (black arrowhead). (C) Quantification of results in (A-B). (D) Mid-gastrula control embryo depicting wildtype *myod1* expression; (E) *rab7* knockdown revealed reduced or absent ventral *myod1* expression in most specimen (black arrowheads). (F) Quantification of results in (D-E). (G) Late gastrula luciferase-based BAR-reporter assay demonstrating inhibition of endogenously induced Wnt reporter activity after co-injection of *rab7*-TB-MO into ventral mesoderm. (H) Quantification of epistatic function of Rab7 and Ctnnb1 in ventro-lateral mesoderm, different manifestations of *myod1* expression in analyses are exemplified below quantification. *Abbreviations as indicated in Fig. 1*

Therefore, we analyzed Wnt-dependent ventro-lateral genes during late gastrulation, to test if *rab7* was required for *wnt8a*-dependent specification of ventral fates. Expression of *myogenic differentiation 1* (*myod1*) and *T-box 6* (*tbx6*) were strongly inhibited or lost in morphants, demonstrating a requirement of *rab7* for ventral mesoderm identity (Fig. 5D-F; Fig. S5C-E). We also injected the BAR reporter in this area to directly quantify the effect of *rab7* inhibition on endogenous Wnt activity. Loss of *rab7* reduced Wnt reporter signals by about 90%, strongly indicating that the loss of marker genes was caused by Wnt pathway inhibition (Fig. 5G). To get further support for this conclusion, we performed epistasis experiments using suboptimal doses of the *rab7*-TB-MO together with a well-established *ctnnb1*(*β-catenin*)-MO (Heasman et al., 2000). While injection of effective doses of *ctnnb1*-MO resulted in loss of *myod1* expression comparable to *rab7* morphants, low-dose injections of either *ctnnb1*-MO or *rab7*-TB-MO both had only minor effects on ventral *myod1* (Fig. 5H and Fig. S5F-I). When both MOs were combined using low doses, *myod1* expression was lost in all double-morphants examined, supporting the conclusion of an epistatic interaction of Rab7 and Ctnnb1 (Fig. 5H and Fig. S5I).

Together, this set of experiments supports the conclusion that *rab7* is necessary for endogenous Wnt8a-mediated specification of ventral and somitic fates during gastrulation, downstream of Wnt receptor activation.

### Rab7 acts epistatically with the endosomal regulator Vps4

In most contexts, Rab7 acts via its well-studied role as a regulator of late endosomal function. However, in some cases it has been shown to perform a cellular role independent of LE, and potentially not in the endo-lysosomal pathway (Guerra and Bucci, 2016). Therefore, we aimed to address this point here as well, namely by testing if other late endosomal regulators known to be required for Wnt pathway function would co-regulate ventral, Wnt-dependent fates in concert with Rab7. We chose two components of the ESCRT machinery (Horner et al., 2018), which have previously been characterized in Wnt signal transduction in *Xenopus*, i.e. *hepatocyte growth factor-regulated tyrosine kinase substrate* (*hrs*) and *vacuolar protein sorting 4 homolog* (*vps4*; Taelman et al., 2010).

Using doses that have previously been shown to block double axis formation, we then knocked down *hrs* in the ventral mesoderm, or overexpressed a dominant-negative version of *vps4* (*dnvps4*; Bishop and Woodman, 2000), to analyze Wnt-dependent mesoderm patterning. In both cases, loss of these late endosomal regulators also caused reduction or loss of *myod1* in the ventral part, demonstrating their necessity for correct patterning (Fig. 6A,B,E and Fig. S6A-C). As this implicated a functional cooperation with Rab7 on LE, we next performed an epistatic analysis to demonstrate interaction. Either low-dose injection of *dnvps4* mRNA, or that of *rab7*-TB-MO caused only minor reduction of *myod1* expression, however, parallel injection of both agents caused strong inhibition of *myod1* in a significant manner (Fig. 6B-E). From these results we conclude that Rab7 participates in Wnt-dependent patterning of the ventral mesoderm as an endosomal regulator together with other effectors of normal LE function.

**Fig. 6.**
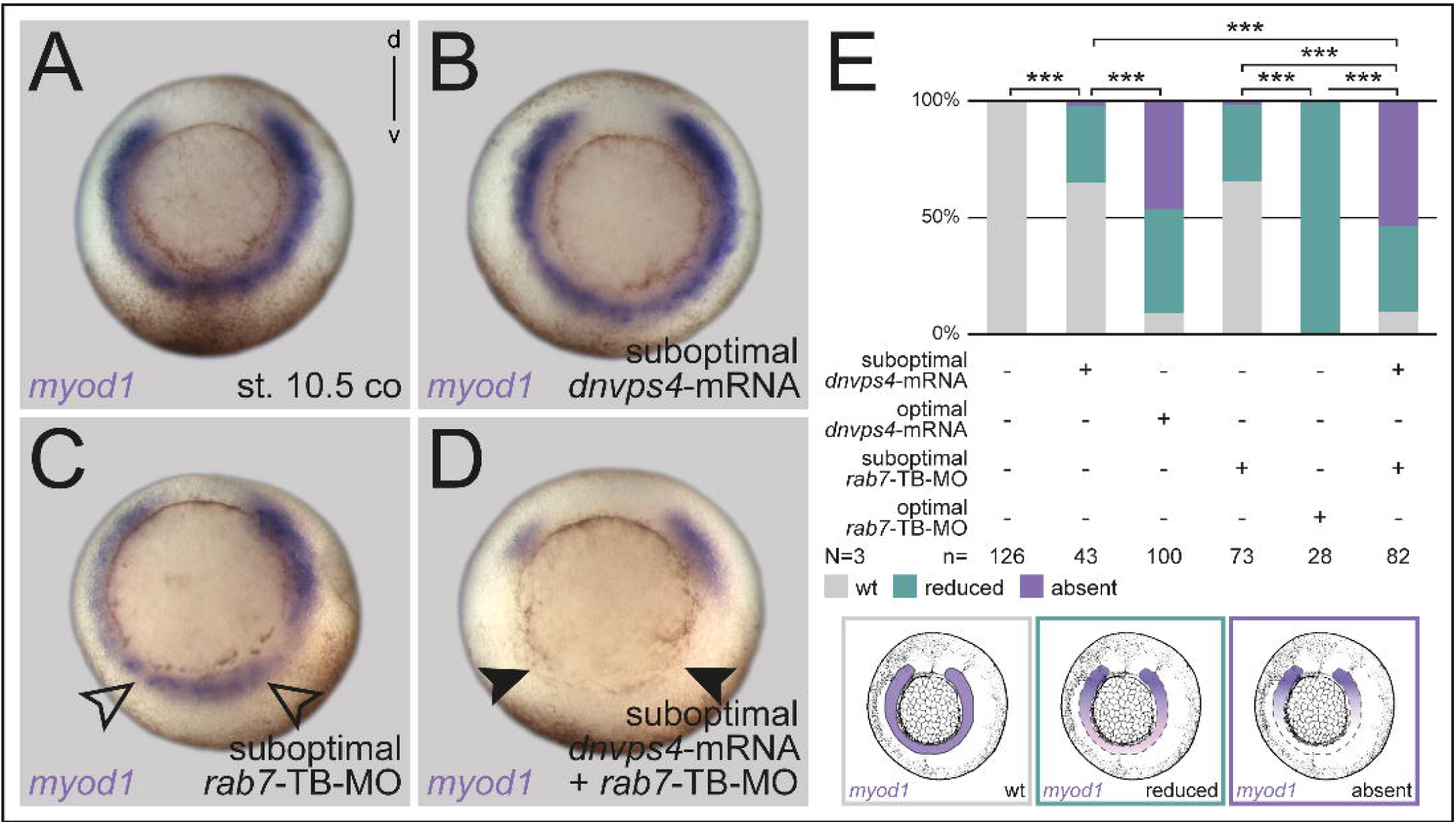
Small GTPase Rab7 acts epistatically with the LE regulator Vps4. (A) Untreated specimen. (B) Injection of *dnvps4* mRNA or (C) *rab7*-TB-MO in suboptimal dose lead to minor reduction of *myod1* expression on ventral side (black outlined arrowheads). (D) Parallel suboptimal injection of *rab7*-TB-MO and *dnvps4* mRNA resulted in absent *myod1* expression (black arrowheads). (E) Quantification of results, different manifestations of *myod1* expression in analysis are exemplified below quantification. *Abbreviations as indicated in Fig. 1*

## Discussion

In this study, we analyzed the *in vivo* role of the small GTPase Rab7 in the frog *Xenopus* with focus on its role in early patterning and regulation of morphogenetic processes during gastrulation. We were able to demonstrate a requirement for Wnt-dependent processes in the early embryo, both, endogenously in the mesoderm, or if exogenously induced. Further, our analyses implicated that Rab7 might participate in further Wnt-dependent, or putatively independent processes.

### *rab7* has distinct activity pattern throughout embryonic development

In developmental studies, genes with housekeeping function are mostly used as molecular tools – developmental expression is rarely included (Kim et al., 2012; Lee and Harland, 2010 for Rab coding examples). Our analysis of *rab7* revealed strikingly dynamic and spatially restricted expression patterns throughout early development, suggesting a tissue- and context-specific requirement. Therefore, we would predict that many housekeeping genes are enriched in specific tissues, and expression of different endosomal regulators (e.g. *hrs* or *vps4*) could reveal distinct *endosomal synexpression groups*, following a concept proposed before (Niehrs and Pollet, 1999). Such analyses could reveal novel roles for endosomal regulation of developmental processes or signaling pathways, in line with common transcriptional regulation, as demonstrated for genes coding endosomal components (Ploper et al., 2015; Sardiello et al., 2009). In the case of *rab7*, strong signals in the neural plate border (Fig. 1F-H), pronephric kidney, and cranial and trunk neural crest (Fig. 1I,J) suggest such possible roles in other Wnt-regulated tissues (Borday et al., 2018; Burstyn-Cohen et al., 2004; Honoré et al., 2003; Pla and Monsoro-Burq, 2018; Villanueva et al., 2002; Zhang et al., 2011).

### Rab7 is required for dorsal mesoderm specification and tissue remodeling during gastrulation

In our approach presented here, we performed loss of function of *rab7* using antisense oligos and CRISPR/Cas9-mediated genome editing. Reflecting the late expression in diverse tissues, mildly affected, or low-dose injected tadpoles displayed shortened AP axes, head and eye defects, and edema formation, the latter putatively due to loss of *rab7* in the pronephric system (cf. Fig. 1 and S1; Wessely and Tran, 2011). Yet, while developing no phenotypes during cleavage and blastula stages, the most prominent developmental defects of *rab7*-deficient embryos became apparent during gastrulation, as a result of both, morphogenetically as a lack of CE and notochord specification on the dorsal side, and ventrally due to a lack of mesoderm specification and subsequent posterior truncations. The lack of very early phenotypes despite the presence of a large supply of maternal *rab7* mRNA in cleavage and blastula stages suggest that this pool might be required mainly after midblastula transition (MBT), i.e. at least partially during gastrulation. This is supported by the fact that the *rab7*-TB-MO was slightly more efficient in causing gastrulation defects than the *rab7*-SB-MO. Yet, some maternal Rab7 protein might mask an early role during cleavages as well.

Interestingly, work in mice strongly supports an evolutionary conserved role of *rab7* in this context (Kawamura et al., 2012). Here, *rab7* KO also prevented gastrulation by interfering with AVE development, resulting in early embryonic death. More importantly, in a recently published follow-up report, the authors demonstrated that this gastrulation phenotype was caused by lack of proper mesoderm formation, recognizable by a failure in correct tissue remodeling and subsequent germ layer formation (Kawamura et al., 2020). These cellular phenotypes are highly reminiscent of our observations in the dorsal mesoderm after loss of *rab7*, where cellular arrangements were disorganized as well, and correct formation of Brachet’s cleft was impaired, i.e. germ layers also failed to separate correctly (cf. Fig. 2 C-M and Fig. S2). Another indication for a conserved role of *rab7* in these processes is the reduction of cell adhesion markers we observed (cf. Fig. 2K-M and Fig. S2A-D). In the mouse, such altered cell adhesion was also observed after loss of *rab7*, paralleling the failed tissue remodeling (Kawamura et al., 2020).

From work in cell culture it is known that Rab7 is required for correct activation and localization of β1-Integrin at the cell membrane in a permissive way, i.e. as a component required for transport towards the membrane (Arjonen et al., 2012; Margiotta et al., 2017). Thus, such a function of Rab7-mediated transport would explain the reduction of β1-Integrin we observed after rab7 inhibition (Fig. 2L,M). Yet, we cannot clearly distinguish whether the reduction of β1-Integrin (and Ctnnb1) in the dorsal mesoderm is a direct result of loss of Rab7-mediated transport of these molecules, or an indirect consequence of the lack of mesodermal specification. The significant down-regulation of *tbxt* in most embryos might argue for the second possibility (Fig. 3P-R), which could explain all observed problems of tissue remodeling, CE and notochord morphogenesis – as Tbxt is a well-known upstream regulator of mesoderm specification and non-canonical Wnt pathways required for CE (Bruce and Winklbauer, 2020; Schulte-Merker and Smith, 1995; Tada and Smith, 2000).

### Rab7 mediates dorsal development independent of the organizer

Strikingly, we did not observe a change in organizer induction as judged by expression of *gsc* or *chd* at early gastrulation. Further, the embryos show no signs of ventralization or dorsalization, neither phenotypically, nor when analyzed for mid-gastrula expression of the DV-specific genes *ventx1* or *chd* (Figs. 3D-F and S3F-K). These results argue against an involvement of *rab7*, first, in early Wnt-mediated Nieuwkoop center/organizer induction (Fagotto et al., 1997; Heasman et al., 1994), second, in TGF-β/Nodal pathway-induced activation of organizer genes, and third, in BMP-mediated ventral development (E.M. De Robertis, 2009). These conclusions are again supported by the recent report showing also no alteration of Nodal or BMP signaling in *rab7* KO mice (Kawamura et al., 2020). For the TGF-β members BMP and Nodal it is known that endocytosis and subsequent translocation of activated receptor complexes to EE is required for intracellular signal transduction, i.e. correct function of Rab5-mediated processes should be a prerequisite of this (Platta and Stenmark, 2011; Sigismund et al., 2012). However, in *Xenopus*, Rab5-mediated endocytosis has been reported neither to be required for long-range signaling, nor for cells generally to respond to activin ligands (i.e. another TGF-β member) during mesendoderm induction (Hagemann et al., 2009). So far, an involvement of Rab7 or LE in general has not been linked to the activation of TGF-β signaling, a finding we also conclude from our *Xenopus* analyses. However, we cannot exclude a role in specific tissues during later development.

In contrast to the wildtype expression of organizer genes, we found that *rab7* was clearly necessary for *myf5, foxj1* and *tbxt*, and partially for correct *nodal3* expression (Figs. 3G-R). These genes are known to depend on active Wnt signaling (although with some uncertainty for *tbxt*), however, it is not fully understood to which extent the maternally or zygotically activated Wnt signals contribute to their activation (Shi et al., 2002; Smith et al., 1995; Stubbs et al., 2008; Vonica and Gumbiner, 2002; Walentek et al., 2013). *nodal3* expression, which was impacted least after *rab7* inhibition, is initiated very early on and is thought to be a direct Wnt target (Glinka et al., 1996; Smith et al., 1995). Therefore, these results might indicate a role for Rab7 only for processes relying (at least partially) on zygotic Wnt activation. In the recently analyzed rab7 KO mouse, gastrulation phenotypes and lack of *tbxt* expression have been demonstrated to be related to mis-regulation of Wnt signaling by upregulating the activity of the Wnt antagonist Dickkopf (Dkk; Kawamura et al., 2020). While this, probably mammal-specific, AVE-to-epiblast crosstalk of Wnt activity modulation cannot easily be transferred to *Xenopus* gastrula embryos, the common Wnt-dependent induction or maintenance of *tbxt* seems to support a conserved general role of *rab7*. Yet, a further confirmation of this Wnt connection comes from another analysis in mouse, where notochord-specific inhibition of Wnt signaling resulted in reduced expression of *noto, shh* and *tbxt*, resulting in posterior truncations as well (Ukita et al., 2009).

### Rab7 is necessary for Wnt activation in *Xenopus* in a context-dependent manner

The unexpected lack of ventralization phenotype contradicted the reported role of LE for Wnt pathway activation, as we expected Rab7 to be also required for maternal Wnt-dependent organizer induction (Taelman et al., 2010; Vinyoles et al., 2014). This was even more puzzling, as we could demonstrate an absolute requirement for exogenously induced activation of Wnt-dependent processes, i.e. double axis assay, Wnt reporter activation, and the restoration of the DV axis after Wnt8a-mediated dorsalization – and some of these processes act also early during cleavage stages (cf. Figs. 4 and S4). The last result exemplified this discrepancy clearly, as exogenously induced Wnt-dependent dorsal fates were blocked after loss of *rab7*, while the endogenous, organizer-induced expression of *chd* and *nog* stayed unaltered (cf. Fig. 4A-D and S4A-D). A compensatory action by the paralogous *rab7b* can be excluded, as it is not present in the zygote (Peshkin et al., 2019; Session et al., 2016; our unpublished data). One possible, straightforward explanation would be that endogenous early Wnt activation does not require ligand-mediated receptor activation and endocytosis, which we used for our experiments with exogenous Wnt pathway induction. This could thus include a mechanism that bypasses the endolysosomal system. However, it has been reported that organizer induction does require autocrine and paracrine Wnt ligand perception (Cha et al., 2008; Kofron et al., 2007; Tao et al., 2005). Another possibility would be that maternal Wnt does not use LE at all. In cell culture and in *Xenopus* embryos, LE have been reported to be necessary to establish a robust Wnt output, i.e. to maintain continuous inhibition of glycogen synthase kinase 3 (GSK3), and thus Wnt pathway activation. However, endogenous organizer induction has not been analyzed in these experiments (Niehrs and Acebron, 2010; Taelman et al., 2010). In fact, we cannot exclude that early dorsal Wnt activation might only rely on ‘fast-acting’, LE-independent mechanisms (like Axin inhibition), which have been suggested to be required for GSK3-inhibition and Wnt activation (cf. Clevers and Nusse, 2012; Li et al., 2012). Alternatively, the *Xenopus* zygote might already contain fully or partially matured LE, as proposed before, but whose functionality we are not able to interfere with using embryonic injections (Dobrowolski and De Robertis, 2011). Finally, there might be simply enough maternally deposited Rab7 protein present, just sufficient to participate in organizer induction. Yet, due to the close temporal proximity of endogenous organizer induction and that mediated by our *wnt8a* mRNA injections, we would question this possibility. Future analyses might reveal the nature of this phenomenon.

In contrast to the rather complex involvement of Rab7 for dorsal Wnt-mediated fates, we could show a requirement for endogenous ventro-lateral mesoderm patterning, as all marker genes (*myf5, tbx6, myod1*) were strongly downregulated after loss of *rab7* (Figs. 3M-O, 5D-F, S5C-E). These genes have all been shown to depend on a ventral source of Wnt, mediated by Wnt8a (Hoppler and Moon, 1998; Hoppler et al., 1996; Kjolby and Harland, 2017; Shi et al., 2002), which was supported by the epistatic effect of *rab7* knockdown with *ctnnb1* knockdown (Figs. 5H and S5F-I). Furthermore, we could also demonstrate that *rab7* knockdown strongly blocked endogenous Wnt target gene activation, as monitored using a ventral mesodermal BAR reporter signal (Fig. 5G). Finally, *ventx1* expression was also reduced after ventral injections (Figs. 3D-F and S4E-G), which is in agreement with a recent report showing such a reduction of *ventx1* after ventral loss of (zygotic) *wnt8a* (Nakamura et al., 2016). From these experiments we conclude that *rab7* participates in ventral mesoderm specification by acting as a necessary factor for Wnt activation. This effect on mesoderm specification was phenocopied by loss of LE-associated ESCRT factors *vps4* and *hrs*, which are known to be required for Wnt pathway activation (Taelman et al., 2010). As we could demonstrate an additive relationship with loss of *rab7* (Fig. 6), we conclude that Rab7 fulfills this role as an endosomal regulator required for correct LE-mediated Wnt transduction (Dobrowolski and De Robertis, 2011; Hikasa and Sokol, 2013).

### The connection of Rab7 to other signaling pathways – beyond Wnt activation

While our findings about a role of Rab7 in *Xenopus* Wnt pathway activation seem clear for exogenously induced and for endogenous ventral mesoderm specification, some results with dorsal marker gene might indicate further, perhaps Wnt-independent roles of Rab7 during gastrulation. While dorsal loss of *rab7* resulted in gastrulation and CE defects, selective downregulation of *shh* seems puzzling (Fig. 2Q-S). Regulation of *shh* in the ventral neural tube is well-analyzed (Dessaud et al., 2008), however, surprisingly little was reported about induction and maintenance of its notochordal expression. Activin was shown to be able to induce *shh* in animal caps, but not endogenously in the mesoderm (Yokota et al., 1995), and we have no evidence for a participation of Rab7 in TGF-β pathways for now. Yet, in the well-studied limb bud, Wnt7a has been shown to be required for induction and/or maintenance of *shh*, offering a potential link to our observations (Parr and McMahon, 1995; Yang et al.). Another interesting aspect also comes from the limb bud, where fibroblast growth factor (FGF) signaling was reported both, to induce and to maintain *shh* expression (Scherz et al., 2004; Vogel et al., 1996; Yang et al.). FGF signaling is known to cooperate with Wnt in different contexts, and both are required additively to induce *myf5* in the somitic mesoderm (e.g. Shi et al., 2002). Thus, if *rab7* also participated in FGF pathway activation in some way, this would explain the strong effect on *myf5* after loss of function – and the differential impact on *nodal3* versus *foxj1* and *tbxt* (Fig. 3G-R). All depend on Wnt signaling, but only *tbxt* and *foxj1* have been shown to be regulated by the FGF pathway dorsally, downstream of Nodal3-induced activation of Fgf receptor 1 (Glinka et al., 1996; Schneider et al., 2019; Smith et al., 1995; Vick et al., 2018; Yokota et al., 2003)!

## Materials and methods

### *Xenopus laevis* care and maintenance

Frogs were purchased from Nasco (901 Janesville Avenue P.O. Box 901 Fort Atkinson). Handling, care and experimental manipulations of animals was approved by the Regional Government Stuttgart, Germany (V349/18ZO ‘Xenopus Embryonen in der Forschung’), according to German regulations and laws (§6, article 1, sentence 2, nr. 4 of the animal protection act). Animals were kept at the appropriate conditions (pH=7.7, 20°C) at a 12 h light cycle in the animal facility of the Institute of Zoology of the University of Hohenheim. Female frogs (4-15 years old) were injected subcutaneously with 300-700 units of human chorionic gonadotropin (hCG; Sigma), depending on weight and age, to induce ovulation. Sperm of male frogs was gained by dissection of the testes that was stored at 4°C in 1x MBSH (Modified Barth’s saline with HEPES) solution. Embryos were staged according to Nieuwkoop and Faber (1994). Only clutches of embryos from healthy females were used for the experiments reported here – provided the early embryonic stages showed normal survival rates as well. Individual embryos from one batch were randomly picked and used either as control or tested specimens. If control groups displayed unusual developmental defects later in development, such clutches were excluded as well, based on empirical judgement.

### Morpholino design and microinjections

The *rab7*-5’UTR-/TB-MO was designed using the sequence of the S-form from the genomic sequence as deposited in gene bank (one mismatched base pair for the L-Form). TB-MO-sequence is 5’-GTCTCCGCTTCCTACCCCTGCCAGC-3’. The *rab7*-SB-MO was designed using the sequence of the L-Form from the genomic sequence as deposited in gene bank (three mismatched base pairs for the S-Form). Splice acceptor site at intron 2 of the *rab7* pre-mRNA is targeted by SB-MO (5’-GCCAACCCTAGAATGGAAGATACAA-3’). Further MOs used in this study were *ctnnb1*- and *hrs*-MO as published (Heasman et al., 2000; Taelman et al., 2010) or a random co-MO as a MO fill up for epistatic analyses. Generally, embryos were injected at the 4-cell stage into both dorsal or ventral blastomeres, respectively, unless indicated differently, using a Harvard Apparatus setup. Drop size was calibrated to 4 nl per injection. Total amounts of injected MOs were: 0.4 pmol *ctnnb1*-MO (suboptimal dose), 0.8 pmol *ctnnb1*-MO (optimal dose), 1.6-2.0 pmol *hrs*-MO, 0.7 pmol *rab7*-TB-MO (suboptimal), 1.0-1.4 pmol *rab7*-TB-MO (optimal), 1.4-4.0 pmol *rab7*-SB-MO.

### mRNA synthesis and microinjections

Plasmids were linearized with NotI and transcribed *in vitro* (Sp6 polymerase) using Ambion message machine kit. Drop size was calibrated to 4 nl per injection. Total amounts of injected mRNA per embryo are as followed: 80 pg *ca-rab7* mRNA, 400 pg *dnvps4* mRNA, 400 pg GFP mRNA, 160 pg *wnt8a* mRNA

### sgRNA design and microinjections

Two single guide RNAs were designed for the *Xenopus laevis rab7a* gene, *rab7*-CRNP (S+L), target sequence: 5’-GGTGATGGTGGATGACAGAT TGG-3’ (on exon 3), and rab7-CRNP (L), target sequence: 5’-GGGACACAGCTGGGCAGGAA AGG-3’ (on exon 4), using the publicly available ‘CRISPscan’ software. sgRNAs were transcribed with the MEGAshortscript T7 Kit from synthetic DNA oligomers and purified with the MEGAclear Transcription Clean-Up Kit (both ThermoFisher).

Oligos for synthesis were: rab7-CRNP (S+L) Forward: 5’-GCAGCTAATACGACTCACTATAGGTGATGGTGGATGACAGATGTTTTAGAGCTAGAAATAGC AAG -3, rab7-CRNP (L) Forward 5’-GCAGCTAATACGACTCACTATAGGGACACAGCTGGGCAGGAAGTTTTAGAGCTAGAAATAG CAAG-3’, and general Reverse_5’-AAAAGCACCGACTCGGTGCCACTTTTTCAAGTTGATAACGGACTAGCCTTATTTTAACTTGCT ATTTCTAGCTCTAAAAC -3’ Embryos were injected with 1 ng Cas9 protein (PNA Bio) and 300 pg sgRNA at 1-cell stage and cultivated at room temperature until desired stage. DNA was isolated from 10 mutant or 5 control embryos. For verification of successful genome editing, isolated DNA was proceeded by RT-PCR and sequenced. Analysis of sequenced DNA was analyzed via Synthego ICE. PCR-Primers for sequencing were the following: *rab7*-CRNP(S+L), L-form (FP 5’-AGCCGTATTTCTTTGTTGTGCCA-3’; RP 5’-ATTCCAGGTGCAGTGAGATGT-3’), *rab7*-CRNP(S+L), S-form (FP 5’-TGAGTGCATTGTGCTGTGTG-3’; RP 5’-CCCCCATTTGAAAACTGAAGAGAG-3’), *rab7*-CRNP(L), (FP 5’ ACGGGAGCAGATTTAATAGGACA-3’; RP 5’-CTTGGACTCGCCTGGATGAG-3’)

### PCR-based verification of efficient intron retention for the SB-MO

For verification of SB-MO, a standard RT-PCR was performed. SB-MO was injected in all 4 blastomeres of 4-cell embryos, cultivated until st. 13, and fixed for RNA isolation. Following PCR primers were used: Forward 5’-CCTCCAGGAATATGCAGGAA-3’; Reverse 5’-CTGCATTGTGACCAATCTGTC-3’

### Luciferase based Wnt assay

For Luciferase based Wnt assay (Promega Dual-Luciferase® Reporter Assay System), embryos were injected into two animal or ventral blastomeres at 4-cell stage (80 pg BAR reporter DNA plus 40 pg Renilla DNA). For exogenously induced reporter activity, embryos were injected of 240 pg *wnt8a* mRNA +/−1.4 pmol *rab7*-TB-MO and cultivated until stage 10.5, then animal caps were dissected and further processed. For endogenous analysis, embryos were co-injected with rab7-TB-MO, cultivated until st. 12, the ventral halves were dissected. Dissection of tissues was performed in 0,1x MBSH. Extracted tissues (10 animal caps or ventral halves) were transferred into lysis buffer (Promega) and incubated for 15 min. Lysates were centrifuged repeatedly for 15 min, then supernatant transferred in triplets onto a 96-well plate for Luciferase analysis by the GloMax Explorer system. Relative luciferase units (RLU in [%]) were calculated by the ratio of Luciferase and Renilla values.

### *In situ* Hybridization

For *in situ* mRNA detection, ISH was performed after fixation in MEMFA for 2-3 h at room temperature and processed following a customized standard protocol (Sive et al., 2000; customization after R. Rupp, personal communication). RNA *in situ* probes were transcribed using SP6 or T7 polymerases.

### Axis induction Assay

Double axis assay was performed by single injection of 0,8 pg *wnt3a* mRNA +/−0,7 pmol *rab7*-TB-MO into one ventral mesodermal blastomere at 4-cell stage as described (Beyer et al., 2012). Embryos were raised until gastrulation or neurula stage. Double axes were scored empirically by second ventral *gsc* expression (early gastrula) and for visible induction of secondary axes (late neurulas).

### Embryo sections

For vibratome sections, embryos were embedded in a glutaraldehyde-crosslinked gelatin-albumin mix and razor blade-sectioned. Hoechst stained vibratome sections were incubated on microscope slide before imaging (1:10000 Hoechst). Bisections of embryos was performed using a razor blade.

### Immunofluorescence Analysis

Co-injected fluorescein dextran (70,000 MW, ThermoFisher, D1822) was used as lineage tracer for dorsal lips in IF analyses. For IF analyses, embryos were fixed in 4% paraformaldehyde for 1 h at RT, followed by 2 washes in 1x PBS-for 15 min each. For staining of animal caps or bisected specimens, embryos were manually dissected horizontally or sagittaly after fixation, transferred to 24 well plates, and washed twice for 15 min in PBST (PBS/0.1% Triton X-100). After blocking for 2h at RT in CAS-Block (1:10 in PBST; ThermoFisher, #008120) blocking reagent was replaced by antibody solution (diluted in CAS-Block) for incubation ON at 4°C. Antibodies used were: β1-integrin (DSHB 8C8-s; 1:70), Ctnnb1 (clone H102; Santa Cruz Biotechnology sc-7199; 1:200), MZ-15 (DSHB; 1:200). Then antibody solution was removed and explants washed twice for 15 min in PBS. Secondary antibodies (ThermoFisher, all 1:1000 in CAS-Block) were incubated for 2 h at RT. Cell borders were visualized using AlexaFluor™Plus 405 Phalloidin (ThermoFisher A30104) overnight at 4°C (1:100 in PBS-). For photo documentation, bisected embryos or caps were transferred onto microscope slides or positioned in low melt agarose on a Petri dish (0.5% low melt agarose in 1x PBS-).

### Photo Documentation

LSM images of IF data were taken with a Zeiss LSM 700 Axioplan2 Imaging microscope and then adjusted using the Zeiss Zen 2012 Blue edition. All other pictures were taken with a Zeiss SteREO Discovery.V12 microscope or an Axioplan2 inverted microscope using AxioVision 4.6. Afterwards Adobe Photoshop CS6 was used for cropping and careful brightness adjustments. All figures were arranged using Adobe Illustrator CS6.

### Statistical analysis

Statistical calculations of marker gene expression patterns were performed using Pearson’s chi-square test (Bonferroni corrected, if required). *=p<0.05, **=p<0.01, ***=p<0.001 were used for all statistical analyses, as well as the declaration N = the number of experiments (i.e. number of biological replicates of batches of embryos from different fertilizations), and n = the number of embryos analyzed (i.e. number of biological replicates of embryos).

## Acknowledgements

We like to thank T. Ott for technical advice and valuable feedback, V. Andre, M. Maerker, J.-L. Plouhinec, A. Schäfer-Kosulja, and S. Vogel for technical assistance, and M. Blum and Lab members for continuous valuable feedback. Further, we want to thank Drs. E. De Robertis, R. Harland, T. Hollemann, C. Kintner, R. Moon, R. Rupp, B. van Deurs, and P. Woodman for sharing constructs.

## References

Arjonen, A., Alanko, J., Veltel, S. and Ivaska, J. (2012). Distinct recycling of active and inactive β1 integrins. Traffic 13, 610–625.

Beyer, T., Danilchik, M., Thumberger, T., Vick, P., Tisler, M., Schneider, I., Bogusch, S., Andre, P., Ulmer, B., Walentek, P., et al. (2012). Serotonin Signaling Is Required for Wnt-Dependent GRP Specification and Leftward Flow in Xenopus. Current Biology 22, 33–39.

Biechele, T. L., Adams, A. M. and Moon, R. T. (2009). Transcription-based reporters of Wnt/beta-catenin signaling. Cold Spring Harbor Protocols 2009, pdb.prot5223.

Bishop, N. and Woodman, P. (2000). ATPase-defective mammalian VPS4 localizes to aberrant endosomes and impairs cholesterol trafficking. Mol. Biol. Cell 11, 227–239.

Borday, C., Parain, K., Thi Tran, H., Vleminckx, K., Perron, M. and Monsoro-Burq, A. H. (2018). An atlas of Wnt activity during embryogenesis in Xenopus tropicalis. PLoS ONE 13, e0193606.

Bruce, A. E. E. and Winklbauer, R. (2020). Brachyury in the gastrula of basal vertebrates. Mechanisms of Development 163, 103625.

Brunt, L. and Scholpp, S. (2018). The function of endocytosis in Wnt signaling. Cell. Mol. Life Sci. 75, 785–795.

Burstyn-Cohen, T., Stanleigh, J., Sela-Donenfeld, D. and Kalcheim, C. (2004). Canonical Wnt activity regulates trunk neural crest delamination linking BMP/noggin signaling with G1/S transition. Development 131, 5327–5339.

Butler, M. T. and Wallingford, J. B. (2017). Planar cell polarity in development and disease. Nat Rev Mol Cell Biol 18, 375–388.

Cha, S. W., Tadjuidje, E., Tao, Q., Wylie, C. and Heasman, J. (2008). Wnt5a and Wnt11 interact in a maternal Dkk1-regulated fashion to activate both canonical and non-canonical signaling in Xenopus axis formation. 135, 3719–3729.

Clevers, H. and Nusse, R. (2012). Wnt/β-catenin signaling and disease. Cell 149, 1192–1205.

Dessaud, E., McMahon, A. P. and Briscoe, J. (2008). Pattern formation in the vertebrate neural tube: a sonic hedgehog morphogen-regulated transcriptional network. Development 135, 2489–2503.

Dobrowolski, R. and De Robertis, E. M. (2011). Endocytic control of growth factor signalling: multivesicular bodies as signalling organelles. Nature Publishing Group 13, 53–60.

E.M. De Robertis (2009). Spemann’s organizer and the self-regulation of embryonic fields. Mechanisms of Development 1–17.

Fagotto, F. (2020). Tissue segregation in the early vertebrate embryo. Seminars in Cell & Developmental Biology.

Fagotto, F., Guger, K. and Gumbiner, B. M. (1997). Induction of the primary dorsalizing center in Xenopus by the Wnt/GSK/beta-catenin signaling pathway, but not by Vg1, Activin or Noggin. Development 124, 453–460.

Fürthauer, M. and González-Gaitán, M. (2009). Endocytic Regulation of Notch Signalling During Development. Traffic 10, 792–802.

Glinka, A., Delius, H., Blumenstock, C. and Niehrs, C. (1996). Combinatorial signalling by Xwnt-11 and Xnr3 in the organizer epithelium. Mechanisms of Development 60, 221–231.

Guerra, F. and Bucci, C. (2016). Multiple Roles of the Small GTPase Rab7. Cells 5, 34.

Hagemann, A. I., Xu, X., Nentwich, O., Hyvonen, M. and Smith, J. C. (2009). Rab5-mediated endocytosis of activin is not required for gene activation or long-range signalling in Xenopus. Development 136, 2803–2813.

Hanson, P. I. and Cashikar, A. (2012). Multivesicular Body Morphogenesis. Annu. Rev. Cell Dev. Biol. 28, 337–362.

Heasman, J., Crawford, A., Goldstone, K., Garner-Hamrick, P., Gumbiner, B., McCrea, P., Kintner, C., Noro, C. Y. and Wylie, C. (1994). Overexpression of cadherins and underexpression of beta-catenin inhibit dorsal mesoderm induction in early Xenopus embryos. Cell 79, 791–803.

Heasman, J., Kofron, M. and Wylie, C. (2000). Beta-catenin signaling activity dissected in the early Xenopus embryo: a novel antisense approach. Developmental Biology 222, 124–134.

Hikasa, H. and Sokol, S. Y. (2013). Wnt Signaling in Vertebrate Axis Specification. Cold Spring Harbor Perspectives in Biology 5, a007955–a007955.

Honoré, S. M., Aybar, M. J. and Mayor, R. (2003). Sox10 is required for the early development of the prospective neural crest in Xenopus embryos. Developmental Biology 260, 79–96.

Hoppler, S. and Moon, R. T. (1998). BMP-2/-4 and Wnt-8 cooperatively pattern the Xenopus mesoderm. Mechanisms of Development 71, 119–129.

Hoppler, S., Brown, J. D. and Moon, R. T. (1996). Expression of a dominant-negative Wnt blocks induction of MyoD in Xenopus embryos. Genes & Development 10, 2805–2817.

Horner, D. S., Pasini, M. E., Beltrame, M., Mastrodonato, V., Morelli, E. and Vaccari, T. (2018). ESCRT genes and regulation of developmental signaling. Seminars in Cell & Developmental Biology 74, 29–39.

Huotari, J. and Helenius, A. (2011). Focus Review Endosome maturation. The EMBO Journal 30, 3481–3500.

Katzmann, D. J., Odorizzi, G. and Emr, S. D. (2002). Receptor downregulation and multivesicular-body sorting. Nat Rev Mol Cell Biol 3, 893–905.

Kawamura, N., Sun-Wada, G.-H., Aoyama, M., Harada, A., Takasuga, S., Sasaki, T. and Wada, Y. (2012). Delivery of endosomes to lysosomes via microautophagy in the visceral endoderm of mouse embryos. Nat Commun 3, 1071–10.

Kawamura, N., Takaoka, K., Hamada, H., Hadjantonakis, A.-K., Sun-Wada, G.-H. and Wada, Y. (2020). Rab7-Mediated Endocytosis Establishes Patterning of Wnt Activity through Inactivation of Dkk Antagonism. CellReports 31, 107733.

Kim, K., Lake, B. B., Haremaki, T., Weinstein, D. C. and Sokol, S. Y. (2012). Rab11 regulates planar polarity and migratory behavior of multiciliated cells in Xenopus embryonic epidermis. Dev. Dyn. 241, 1385–1395.

Kjolby, R. A. S. and Harland, R. M. (2017). Genome-wide identification of Wnt/β-catenin transcriptional targets during Xenopus gastrulation. Developmental Biology 426, 165–175.

Kofron, M., Birsoy, B., Houston, D., Tao, Q., Wylie, C. and Heasman, J. (2007). Wnt11/β-catenin signaling in both oocytes and early embryos acts through LRP6-mediated regulation of axin. Development 134, 503–513.

Lee, J.-Y. and Harland, R. M. (2010). Endocytosis Is Required for Efficient Apical Constriction during Xenopus Gastrulation. Current Biology 20, 253–258.

Li, V. S. W., Ng, S. S., Boersema, P. J., Low, T. Y., Karthaus, W. R., Gerlach, J. P., Mohammed, S., Heck, A. J. R., Maurice, M. M., Mahmoudi, T., et al. (2012). Wnt Signaling through Inhibition of &beta;-Catenin Degradation in an Intact Axin1 Complex. Cell 149, 1245–1256.

Margiotta, A., Progida, C., Bakke, O. and Bucci, C. (2017). Rab7a regulates cell migration through Rac1 and vimentin. BBA – Molecular Cell Research 1864, 367–381.

Nakamura, Y., de Paiva Alves, E., Veenstra, G. J. C. and Hoppler, S. (2016). Tissue- and stage-specific Wnt target gene expression is controlled subsequent to β-catenin recruitment to cis-regulatory modules. Development 143, 1914–1925.

Niehrs, C. and Acebron, S. P. (2010). Wnt signaling: multivesicular bodies hold GSK3 captive. Cell 143, 1044–1046.

Niehrs, C. and Pollet, N. (1999). Synexpression groups in eukaryotes. Nature 402, 483–487.

Nieuwkoop, P. D. and Faber, J. (1994). Normal Table of Xenopus laevis (Garland, New York).

Parr, B. A. and McMahon, A. P. (1995). Dorsalizing signal Wnt-7a required for normal polarity of D–V and A–P axes of mouse limb. Nature 374, 350–353.

Peshkin, L., Lukyanov, A., Kalocsay, M., Gage, R. M., Wang, D., Pells, T. J., Karimi, K., Vize, P. D., Wühr, M. and Kirschner, M. W. (2019). The protein repertoire in early vertebrate embryogenesis. Preprint, bioRxiv 1865, 571174.

Pla, P. and Monsoro-Burq, A. H. (2018). The neural border: Induction, specification and maturation of the territory that generates neural crest cells. Developmental Biology 444 Suppl 1, S36–S46.

Platta, H. W. and Stenmark, H. (2011). Endocytosis and signaling. Current Opinion in Cell Biology 23, 393–403.

Ploper, D. and De Robertis, E. M. (2015). The MITF family of transcription factors: Role in endolysosomal biogenesis, Wnt signaling, and oncogenesis. Pharmacol. Res. 99, 36–43.

Ploper, D., Taelman, V. F., Robert, L., Perez, B. S., Titz, B., Chen, H.-W., Graeber, T. G., Euw, von E., Ribas, A. and De Robertis, E. M. (2015). MITF drives endolysosomal biogenesis and potentiates Wnt signaling in melanoma cells. Proc. Natl. Acad. Sci. U.S.A. 112, E420–9.

Sardiello, M., Palmieri, M., di Ronza, A., Medina, D. L., Valenza, M., Gennarino, V. A., Di Malta, C., Donaudy, F., Embrione, V., Polishchuk, R. S., et al. (2009). A Gene Network Regulating Lysosomal Biogenesis and Function. Science 325, 473–477.

Scherz, P. J., Harfe, B. D., McMahon, A. P. and Tabin, C. J. (2004). The Limb Bud Shh-Fgf Feedback Loop Is Terminated by Expansion of Former ZPA Cells. Science 305, 396–399.

Schneider, I., Kreis, J., Schweickert, A., Blum, M. and Vick, P. (2019). A dual function of FGF signaling in Xenopus left-right axis formation. Development 146, dev173575.

Schulte-Merker, S. and Smith, J. C. (1995). Mesoderm formation in response to Brachyury requires FGF signalling. Current Biology 5, 62–67.

Session, A. M., Uno, Y., Kwon, T., Chapman, J. A., Toyoda, A., Takahashi, S., Fukui, A., Hikosaka, A., Suzuki, A., Kondo, M., et al. (2016). Genome evolution in the allotetraploid frog Xenopus laevis. Nature 538, 336–343.

Shi, D.-L., Bourdelas, A., Umbhauer, M. and Boucaut, J.-C. (2002). Zygotic Wnt/beta-catenin signaling preferentially regulates the expression of Myf5 gene in the mesoderm of Xenopus. Developmental Biology 245, 124–135.

Sigismund, S., Confalonieri, S., Ciliberto, A., Polo, S., Scita, G. and Di Fiore, P. P. (2012). Endocytosis and signaling: cell logistics shape the eukaryotic cell plan. Physiological Reviews 92, 273–366.

Sive, H. L., Grainger, R. M. and Harland, R. M. (2000). Early Development of Xenopus Laevis. CSHL Press.

Smith, W. C. and Harland, R. M. (1991). Injected Xwnt-8 RNA acts early in Xenopus embryos to promote formation of a vegetal dorsalizing center. Cell 67, 753–765.

Smith, W. C., McKendry, R., Ribisi, S. and Harland, R. M. (1995). A nodal-related gene defines a physical and functional domain within the Spemann organizer. Cell 82, 37–46.

Sokol, S., Christian, J. L., Moon, R. T. and Melton, D. A. (1991). Injected Wnt RNA induces a complete body axis in Xenopus embryos. Cell 67, 741–752.

Stenmark, H. (2009). Rab GTPases as coordinators of vesicle traffic. Nature Publishing Group 10, 513–525.

Stower, M. J. and Srinivas, S. (2014). Heading forwards: anterior visceral endoderm migration in patterning the mouse embryo. Philos. Trans. R. Soc. Lond., B, Biol. Sci. 369, 20130546.

Stubbs, J. L., Oishi, I., Izpisúa-Belmonte, J. C. and Kintner, C. (2008). The forkhead protein Foxj1 specifies node-like cilia in Xenopus and zebrafish embryos. Nat Genet 40, 1454–1460.

Tada, M. and Smith, J. C. (2000). Xwnt11 is a target of Xenopus Brachyury: regulation of gastrulation movements via Dishevelled, but not through the canonical Wnt pathway. Development 127, 2227–2238.

Taelman, V. F., Dobrowolski, R., Plouhinec, J.-L., Fuentealba, L. C., Vorwald, P. P., Gumper, I., Sabatini, D. D. and De Robertis, E. M. (2010). Wnt Signaling Requires Sequestration of Glycogen Synthase Kinase 3 inside Multivesicular Endosomes. Cell 143, 1136–1148.

Tao, Q., Yokota, C., Puck, H., Kofron, M., Birsoy, B., Yan, D., Asashima, M., Wylie, C. C., Lin, X. and Heasman, J. (2005). Maternal Wnt11 Activates the Canonical Wnt Signaling Pathway Required for Axis Formation in Xenopus Embryos. Cell 120, 857–871.

Teis, D., Wunderlich, W. and Huber, L. A. (2002). Localization of the MP1-MAPK scaffold complex to endosomes is mediated by p14 and required for signal transduction. Developmental Cell 3, 803–814.

Ukita, K., Hirahara, S., Oshima, N., Imuta, Y., Yoshimoto, A., Jang, C.-W., Oginuma, M., Saga, Y., Behringer, R. R., Kondoh, H., et al. (2009). Wnt signaling maintains the notochord fate for progenitor cells and supports the posterior extension of the notochord. Mechanisms of Development 126, 791–803.

Vick, P., Kreis, J., Schneider, I., Tingler, M., Getwan, M., Thumberger, T., Beyer, T., Schweickert, A. and Blum, M. (2018). An Early Function of Polycystin-2 for Left-Right Organizer Induction in Xenopus. iScience 2, 76–85.

Villanueva, S., Glavic, A., Ruiz, P. and Mayor, R. (2002). Posteriorization by FGF, Wnt, and retinoic acid is required for neural crest induction. Developmental Biology 241, 289–301.

Vinyoles, M., Del Valle-Pérez, B., Curto, J., Viñas-Castells, R., Alba-Castellón, L., García de Herreros, A. and Duñach, M. (2014). Multivesicular GSK3 sequestration upon Wnt signaling is controlled by p120-catenin/cadherin interaction with LRP5/6. Molecular Cell 53, 444–457.

Vogel, A., Rodriguez, C. and Izpisúa Belmonte, J. C. (1996). Involvement of FGF-8 in initiation, outgrowth and patterning of the vertebrate limb. Development 122, 1737–1750.

Vonica, A. and Gumbiner, B. M. (2002). Zygotic Wnt activity is required for Brachyury expression in the early Xenopus laevis embryo. Developmental Biology 250, 112–127.

Walentek, P., Schneider, I., Schweickert, A. and Blum, M. (2013). Wnt11b Is Involved in Cilia-Mediated Symmetry Breakage during Xenopus Left-Right Development. PLoS ONE 8, e73646–9.

Wessely, O. and Tran, U. (2011). Xenopus pronephros development--past, present, and future. Pediatr Nephrol 26, 1545–1551.

Yang, Y., Cell, L. N. 1995 Interaction between the signaling molecules WNT7a and SHH during vertebrate limb development: dorsal signals regulate anteroposterior patterning. Cell 80, 939–947

Yokota, C., Kofron, M., Zuck, M., Houston, D. W., Isaacs, H., Asashima, M., Wylie, C. C. and Heasman, J. (2003). A novel role for a nodal-related protein; Xnr3 regulates convergent extension movements via the FGF receptor. Development 130, 2199–2212.

Yokota, C., Mukasa, T., Higashi, M., Odaka, A., Muroya, K., Uchiyama, H., Eto, Y., Asashima, M. and Momoi, T. (1995). Activin Induces the Expression of the Xenopus Homolog of Sonic hedgehog during Mesoderm Formation in Xenopus Explants. Biochem. Biophys. Res. Commun. 207, 1–7.

Zhang, B., Tran, U. and Wessely, O. (2011). Expression of Wnt signaling components during Xenopus pronephros development. PLoS ONE 6, e26533.

